# *Ceratopteris richardii* U6 promoters and plants expressing Cas9 endonuclease as tools for efficient genome editing in a fern

**DOI:** 10.1101/2025.01.12.632625

**Authors:** Robin Schulz, Günter Theißen

## Abstract

Ferns and their allies (monilophytes) represent the second most species-rich group of land plants and are of considerable ecological importance. As a sister group of seed plants (including flowering plants) they are also of great evolutionary interest. Compared to flowering plants, however, much less is known about the developmental and molecular biology of ferns. Among the most important reasons have been the huge genome sizes of ferns and technical obstacles such as the lack of an efficient transformation system. In recent years the situation has improved considerably, however. For the fern model system *Ceratopteris richardii* a whole genome sequence has been published, and an efficient transformation system has been developed. To further facilitate studies on fern biology we aim at simplifying genome engineering of *C. richardii* with the CRISPR-Cas9 system. We report *C. richardii* plants that express Cas9 nuclease under control of the strong CaMV 35S promoter. For efficient expression of single guide RNA (sgRNA) by RNA polymerase III we identified *C. richardii* U6 promoters. These technical improvements may foster many fields of fern physiology, development and evolution.

## Introduction

Monilophytes are a clade of highly diverse land plants (embryophytes). In addition to the large group of ferns *sensu stricto* (with about 10.500 extant species) they also include less-well known and less-species-rich groups such as horsetails (equisetophytes) and whisk ferns (psilophytes) (Pryer et al., 2001). Ferns are of great importance in diverse ecosystems as colonizers, or keystone or invasive species (Marchant et al., 2022). Extant monilophytes represent the sister group of extant seed plants (spermatophytes, that is gymnosperms and angiosperms (flowering plants)) (One Thousand Plant Transcriptomes Initiative, 2019). They are hence of considerable evolutionary interest (Pryer et al., 2001). Therefore, monilophyte phylogeny has been studied in detail to better understand the diversification of both ferns and angiosperms (Schneider et al., 2004). As sister group of spermatophytes monilophytes are essential in studies aiming at a better understanding of some intriguing evolutionary novelties of seed plants such as ovules and seeds.

In comparison to angiosperms, however, only few studies have focused on the molecular biology of ferns (or any other monilophyte). Among the reasons that may have contributed to this neglect are quite certainly their huge genomes, very high chromosome numbers, and, until recently, lack of transformation systems, to mention just a few.

One fern species, however, *Ceratopteris richardii* (also known as “C-fern”), has been established as an upcoming model system already decades ago (Hickok et al., 1995; Eberle et al., 1995). For example, sex-determination mechanisms were studied using classical mutagenesis (Eberle et al., 1995; Banks, 1994, 1999). However, progress was quite slow due to the lack of efficient tools that are available for angiosperm model plants such as *Arabidopsis thaliana*. In recent years, however, an efficient transformation system has been established (Plackett et al., 2014) and a whole genome sequence has been reported (Marchant et al., 2022). Both developments may considerably speed-up fern molecular biology in the near future.

Fern research might be additionally boosted by the establishment of efficient genome editing methods, such as the CRISPR-Cas9 system. It uses RNA-programmable components that originated from type II CRISPR-Cas systems (Ran et al., 2013; Doudna and Charpentier, 2014). These systems provide eubacteria and archaea with adaptive immunity to plasmids and viruses. The CRISPR-associated (Cas) protein Cas9 is an endonuclease that harnesses a guide sequence with an RNA duplex, trascrRNA:crRNA, to form base pairs with DNA target sequences. This enables Cas9 to introduce a site-specific double-strand break in the target DNA. Repair of these lesions in the cell by non-homologous end joining (NHEJ) or homology-directed repair (HDR) may lead to mutations, such as the insertion or deletion of a few nucleotides (Doudna and Charpentier, 2014). If this happens in protein-coding sequences, it often leads to translational frameshift mutations and premature stop codons. This causes truncated proteins during translation or even nonsense-mediated decay (NMD) of the mRNA. Therefore, targeting protein coding regions by the CRISPR-Cas9 system frequently leads to knock-out mutations and thus provides an efficient way of studying gene function by reverse genetics.

The dual tracrRNA:crRNA was engineered as a single guide RNA (sgRNA) that retained two crucial features: a sequence at the 5’ side that determines DNA target site base-pairing and a duplex RNA structure at the 3’ side of the RNA that binds to Cas9 (Ran et al., 2013; Doudna and Charpentier, 2014). This way a simple two-component system was established in which changes in the guide sequence of the sgRNA program the Cas9 nuclease to target any DNA of interest.

It has recently been reported that genome editing employing the CRISPR-Cas9 system could be successfully applied in *C. richardii* to mutate *CrPINMa*, a gene encoding an auxin efflux transporter (Xiang and Li, 2024). However, the genome editing approach used by Xiang and Li (2024) appears quite cumbersome still. To further simplify genome editing in a fern we report here about *C. richardii* plants that express Cas9 constitutively under control of the strong CaMV 35S promoter, and about *C. richardii* U6 promoters to drive sgRNA expression.

## Materials and methods

### Plant materials and growth conditions

Cultivation of *C. richardii* plants (strain Hn-n) was done as generally described (Schulz and Theißen, 2025).

### Identification of endogenous U6-promoters for expression of sgRNA

To find a suitable RNA polymerase III promoter to drive the expression of sgRNA we started with the popular *Arabidopsis thaliana* U6 promoter AtU6-26 (AT3G07265; Zhang et al., 2016) as a reference to find potential *Ceratopteris* specific U6 promoters. We performed a BLAST search (nucleotide vs. nucleotide) with the sequence of AtU6-26 as query against the genome of *Arabidopsis thaliana* using the databases TAIR10 and Araport11 to find the corresponding transcripts in *A. thaliana*. Those transcripts were then used for a BLAST search (nucleotide vs. nucleotide) on phytozome-next.jgi.doe.gov against the genome of *C. richardii* using the database *C. richardii* 2.1 to find *Ceratopteris* specific transcripts driven by a U6 promoter. Genomic regions of 1 kb upstream of 50 selected transcripts were screened for the presence of the two well conserved essential promotor elements in plants known as upstream sequence elements (USE) with the sequence of 5’-RTCCCACATCG-3’ and TATA-box (Filipowicz et al., 1990). We took care that there were no recognition sites for the restriction enzymes *ECO31I* and *BpiI*, which are used in the golden gate cloning. The promoters were named CrU6 followed by minus and number of the candidate.

### Generation of constructs for expression of Cas9 and sgRNA

On the following every PCR was done using Phusion polymerase. The construct pB-35S∷*Cas9* was created by cloning the CDS of *Cas9*, codon optimized for *A. thaliana*, as a *Bam*HI-*Xba*I fragment into modified pBOMBER (Plackett et al., 2014). Template for the *Cas9* was the vector pDE-Cas9 (Fauser et al., 2014). The recombination sites for *Bam*HI and *Xba*I were added to the CDS of *Cas9* by mutagenic PCR. Mutagenic primers were as follows: forward: 5’-CGGGGATCCATGGATAAGAAGTAC-3’; reverse: 5’-AGCTCTAGAATCACCACCGAGC-3’. The PCR program was first 98 °C for 2 min, then 10 cycles with 98 °C for 15 s, annealing at 62.5 °C for 15 s and elongation at 72 °C for 3 min, and then 20 cycles with the annealing changed to 65.2 °C, then 72 °C for 10 min. Selection of *E. coli* was for both plasmids pBOMBER and pDE-Cas9 on 50 μg/mL [w/v] spectinomycin (ThermoFisher, Germany), and therefore the *Cas9* gene was first cloned into pJET1.2 (ThermoFisher, Germany), named pJET-Cas9, to change the resistance marker to 100 μg/mL [w/v] ampicillin (ThermoFisher, Germany). The *GUS* gene of the original pBOMBER plasmid was cut out using *Bam*HI-*Xba*I (FastDigest, ThermoFisher, Germany) and the linear plasmid was isolated with gel purification (GeneJET Gel Extraction Kit, ThermoFisher, Germany). *Cas9* was cut out of pJET-Cas9 with *BamHI* and *XbaI*, purified on a gel and ligated into the *Bam*HI-*Xba*I overhangs of the linear pBOMBER using T4 DNA ligase (ThermoFisher, Germany). The construct was called pB-Cas9 and uses the CaMV 35S promoter for the expression of *Cas9* with OCS terminator. The selection was controlled by *hygromycin phosphotransferase* (*hpt*) for plant selection on hygromycin B.

Multiple constructs for the expression of sgRNA were built by golden gate cloning. The plasmid pGGZ001 (Lampropoulos et al., 2013) was used as the backbone. The constructs pGGZ-CrU6 had a CrU6 promoter candidate, *BpiI* recognition site fused to sgRNA for integration of protospacer and *aminoglycoside phosphotransferase* (*APH*) gene for plant selection on G418. The CrU6 promoter candidate was amplified with mutagenic PCR using genomic DNA of *C. richardii* as template. The mutagenic primers are listed in Table 1. The PCR program was first 98 °C for 2 min, then 10 cycles with 98 °C for 15 s, annealing with temperature A (listed in Table 1) for 15 s and elongation at 72 °C for 1 min., and then 20 cycles with the annealing changed to B (listed in Table 1), then 72 °C for 10 min. The PCR product was first cloned into pJET1.2 which was then used as template for PCR with 30 cycles only using annealing temperature B to generate a product without artefacts. The *BpiI* recognition site fused to sgRNA was amplified with PCR using the plasmid pEN-Chimera (Fauser et al., 2014) and the mutagenic primers (forward: 5’-GGTCTCACGGTCTTCG-3’; reverse: 5’-GGTCTCGTAGTAAAAAAAGCACC-3’). The PCR program was first 98 °C for 2 min, then 10 cycles with 98 °C for 15 s, annealing at 45 °C for 15 s and elongation at 72 °C for 10 s, and then 20 cycles with the annealing changed to 59 °C, then 72 °C for 10 min. The *APH* gene is located on the donor plasmid pGGF007 (Lampropoulos et al., 2013). The PCR products were first purified on a column (GeneJET PCR Purification Kit, ThermoFisher, Germany) and together with pGGZ001 and pGGF007 used in golden gate cloning with *ECO31I* (FastDigest, ThermoFisher, Germany).

**Table 1:**
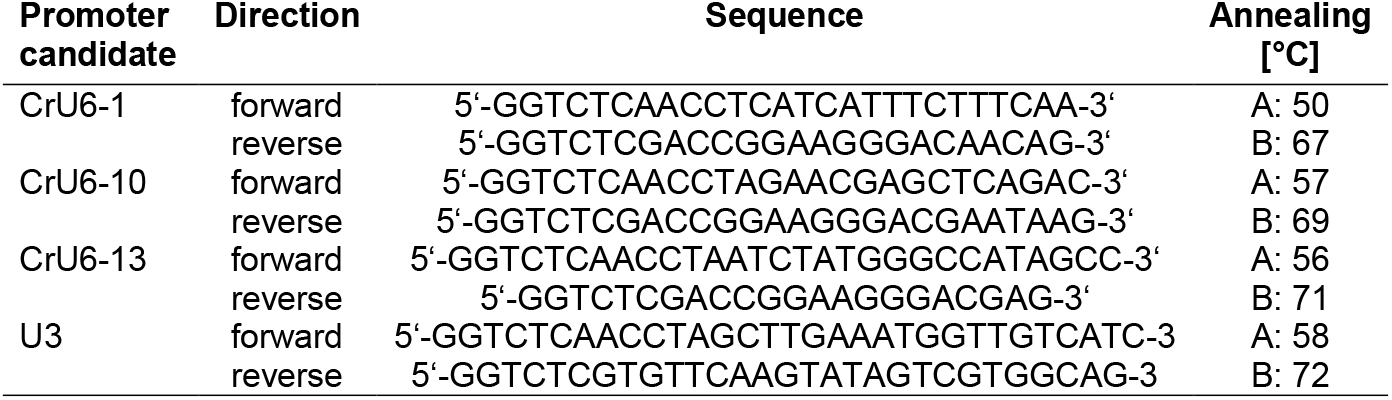
Primer pairs for mutagenic PCR using Phusion polymerase on genomic DNA of *Ceratopteris richardii* to generate CrU6 promoter candidates.

We also build the construct pGGZ-U3 containing a U3 promoter used by Xiang and Li (2024) instead of CrU6. It includes a tRNA (NCBI, AJ506163.1) upstream to the sgRNA. PCR for the U3 promoter used genomic DNA of *C. richardii* as template and the same program as for CrU6 with annealing temperature A (listed in Table 1) for the first 10 cycles and annealing temperature B (listed in Table 1) for the following 20 cycles. The tRNA sequence was acquired (ThermoFisher, Germany) as oligonucleotides (forward: 5’-GGTCTCAAACAAAGCACCAGTGGTCTAGTGGTAGAATAGTACCCTGCCACGGTACAGACCCGGGTTCGATTCCCGGCTGGTGCACGGTCGAGACC-3’; reverse: 5’-GGTCTCGACCGTGCACCAGCCGGGAATCGAACCCGGGTCTGTACCGTGGCAGGGTACTATTCTACCACTAGACCACTGGTGCTTTGTTTGAGACC-3’) containing the *ECO31I* sites and overhangs for golden gate cloning. The volume of each DNA (PCR and plasmid) going into the golden gate reaction was calculated with the formula: 20 (fmol) × size (bp)/concentration (ng/μl)/1520. The program for the golden gate reaction was 30 cycles with restriction digestion at 37 °C for 5 min and ligation at 16 °C for 10 min, then inactivation of *ECO31I* by 60 °C for 5 min and incubated overnight at 4 °C. The correct sequence of all constructs was tested by Sanger sequencing (Macrogen Europe, Netherlands).

An alternative version of pGGZ-CrU6 was build where tRNA-derived DNA sequence was integrated upstream of the sgRNA-derived DNA sequence using pGGZ-U3 as template. The primer pair was: forward: 5’-GGTCTCACTTCAACAAAGCACCAGTGGTC-3’; reverse: 5’-GGTCTCGAAACAGGTCTTCTCGAAGACCG-3’. The PCR used a 2-step program starting with 98 °C for 2 min, then 30 cycles with 98 °C for 15 s, annealing / elongation at 72 °C for 20 s, then elongation at 72 °C for 10 min. The plasmids pGGZ-CrU6-1, pGGZ-CrU6-10 and pGGZ-CrU6-13 were digested with *BpiI* (ThermoFisher, Germany) at 37 °C for 2 h and gel purified. The PCR product for tRNA was digested with *ECO31I* and ligated into the linear pGGZ-CrU6 plasmids.

### Generation of transgenic lines

Generation of callus, transformation and regeneration of callus into sporophytes was performed under aseptic conditions. Transformation was done by microparticle bombardment essentially as described by Plackett et al. (2014). Exceptions are that only callus freshly grown from sporophytes was used for transformation and the callus was selected on 20 mg/L G418 (ThermoFisher, Germany).

### Extraction of total genomic DNA

Total genomic DNA (gDNA) was extracted using amodified cetyltrimethylammonium bromide (CTAB) method with phenol-chloroform. 200 mg freshly frozen fronds (juvenile and fertile) were ground to dust with liquid nitrogen. 1 mL 2 % [w/v] CTAB pH 8.0 (Roth, Germany) combined with 10 μL 100 % ß-mercaptoethanol (Roth, Germany) were heated at 65 °C for 20 min and added to the frozen sample, homogenized by shaking and incubated at 65 °C for 20 min. After centrifugation for 30 min at 16,000 g and 16 °C the upper phase was taken und mixed with 500 μL phenol-chloroform [1:1] (Roth, Germany) and centrifuged for 30 min at 16,000 g and 16 °C. The upper phase was taken and mixed with 0.1 volume 3 M [w/v] NaOAc pH 5.2 (Roth, Germany) and 0.7 volume [v/v] 100 % isopropanol (Roth, Germany). After incubation at room temperature for 60 min centrifugation for 1 h at 16.000 g and 4 °C the liquid was removed with a pipette and the pellet washed with 800 μL 70 % [v/v] ice-cold ethanol (Roth, Germany). The tube was incubated at 4 °C for 15 min, then centrifuged for 15 min. at 16,000 g and 4 °C. All liquid was removed carefully with a pipette and the pellet was dried on the bench at room temperature for 1 h. The DNA was resolved by adding 50 μL Tris-EDTA (TE-buffer) pH 8.0 supplemented with 20 μg/μL RNaseA (ThermoFisher, Germany) and incubated at 65 °C for 1 h, then stored at 4 °C.

### Extraction of total RNA

Total RNA was extracted from freshly frozen fronds (juvenile and fertile) using the RNeasy Kit, Mini, as described by the supplier (Qiagen, Germany). The first centrifugation step was repeated until all visible cell debris from the sample had been removed. The RNA pellet was resolved in 40 μL nuclease free water.

### Identification of transgenic mutants

Identification of mutant plants containing transgenic *Cas9* was done with PCR on genomic DNA and expression of *Cas9* was tested with PCR on complementary DNA (cDNA). The PCR contains the full length of *Cas9* utilizing Phusion polymerase and the primer pair: forward: 5’-ATGGATAAGAAGTACTCTATCGGAC-3’; reverse: 5’-ATCACCACCGAGCTGTGA-3’. The PCR program was first 98°C for 2 min, then 30 cycles with 98 °C for 15 s, annealing at 62.2°C for 15 s and elongation at 72°C for 3 min, then 72°C for 10 min. The PCR product was gel purified, cloned into pJET1.2 and then Sanger sequenced (Macrogen Europe, Netherlands).

### Phenotypic characterization of Cas9 expressing mutants

The major stages of the life-cycle of the *Cas9* expressing mutants in comparison to the wild-type (as described in detail by Conway and Di Stilio, 2019) was documented to determine as to whether the expression of *Cas9* by itself has an obvious impact on the phenotype.

## Results

Gene editing with the CRISPR-Cas9 system (Ran et al., 2013; Doudna and Charpntier, 2014) is optimally using a sgRNA which contains a ∼20 nuncleotide long protospacer at the 5’ end that is complementary to one strand of the target gene DNA sequence, followed by a 76 nt sequence and ending with six thymine as terminator signal. The sgRNA is not translated into a protein. It needs to fold with three hairpins and the 5’ end should not be able to bind to the 3’ end. This structure then binds to the *Cas9* protein and forms a functional endonuclease complex. An important difference is that the *Cas9* gene is transcribed by a DNA-dependant RNA polymerase II (RNAP II) promoter while sgRNA without extensive modification needs a DNA dependant RNA polymerase III (RNAP III) promoter. Previous research has shown that CaMV 35S is a viable promoter for RNAP II. The common promoter to drive RNAP III in plants is the U6 promoter. A characteristic feature of the U6 promoter is that the function is highly species specific. So far, no working U6 promoter for *Ceratopteris* has been reported. We have started by testing the *Arabidopsis thaliana* U6 promoter AtU6-26 that is established for multiple dicot plants (Li et al., 2007) but could not find any sgRNA transcripts (data not shown). We searched, therefore, for endogenous U6 promoters instead and used AtU6-26 as starting point.

After a BLAST search using the AtU6-26 transcript as query against the genome of *Ceratopteris* and filtering the results we found seven novel candidates for *Ceratopteris* specific U6 promoters. Those promoter candidates were named CrU6-1, CrU6-3, CrU6-4, CrU6-10, CrU6-13, CrU6-19 and CrU6-20 (Tab. 1). It is noticeable that most found U6 promoter candidates are located on chromosome 13.

**Table 1:** Promoter candidates of *Ceratopteris richardii* and their position of the genome based on the database phytozome.org and *Ceratopteris richardii* v2.1 reference genome.

| Promoter candidate | Position |
| --- | --- |
| CrU6-1 | Chr13:25175385..25176378 (- strand) |
| CrU6-3 | Chr13:25169205..25170104 (- strand) |
| CrU6-4 | scaffold_1368:81380..82384 (+ strand) |
| CrU6-10 | Chr12:127751051..127752083 (- strand) |
| CrU6-13 | scaffold_1368:90555..91044 (+ strand) |
| CrU6-19 | Chr13:25145175..25146192 (- strand) |
| CrU6-20 | Chr13:25181563..25182580 (- strand) |

Important elements for the function of RNAP II and RNAP III promoters are the upstream sequence element (USE) and TATA box (Fig. 1). The USE is reported to be a highly conserved element in all plants yet in *C. richardii* it has the sequence 5’-AAACCCACATGTG-3’ instead of 5’-RTCCCACATCG-3’ of *Arabidopsis thaliana* (Filipowicz et al., 1990). The TATA box is a basal promoter element which is highly conserved in its position (Wang and Stumph, 1995). The orientation of the TATA box is important to determine whether it is driving RNAP II or RNAP III (Wang and Stumph, 1995). The AtU6-26 promoter of *Arabidopsis* has the reverse TATA box with 5’-TTTATAT-3’. *C. richardii* showed a high conservation of the TATA box with a reverse sequence of 5’-TACATAT-3’. The transcription starting site is 25 to 30 bp downstream of the TATA box (O’Shea-Greenfield and Smale, 1992). The transcriptions starting site (TSS) in both *A. thaliana* and *C. richardii* is a guanine except for CrU6-3 and CrU6-19 which have a cytosine. So far we started testing candidates CrU6-1 and CrU6-10.

**Figure 1:**
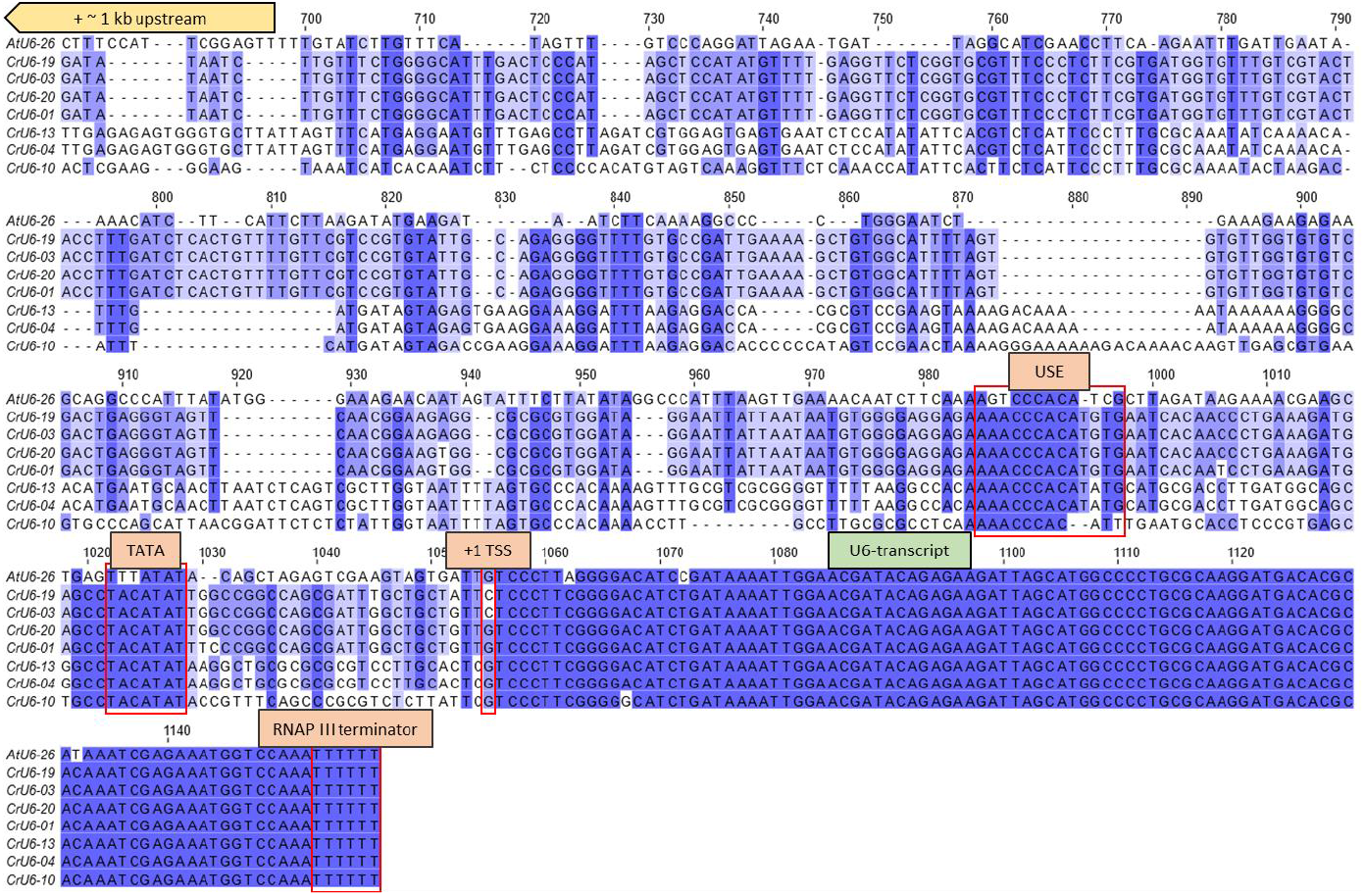
Multiple sequence alignment of candidates of endogenous *Ceratopteris richardii* CrU6 promoters. Around 1 kb upstream of the U6 transcripts were taken as candidates. The conserved elements upstream sequence element (USE) and TATA box show differences between *Ceratopteris richardii* and *Arabidopsis thaliana*. The transcription starting site (TSS) start for both with guanine except for candidate CrU6-03 and CrU6-19 where it starts with cytosine.

Generating constructs for CRISPR-Cas9 is quite difficult and labour intensive. A classical method for generating a CRISPR-Cas9 construct is via golden gate cloning or similar methods. It requires a multitude of different genetic element for the expression of both *Cas9* and sgRNA as well a selection marker (Fig. 2B). At least if done for the first time all individual elements need to be acquired to serve as templates for cloning. The accumulation of all elements increases the size of the plasmid by a large amount (transgenic cassette: ∼12 kb) and therefore also causes a higher probability of shearing when used with a gene gun.

**Figure 2:**
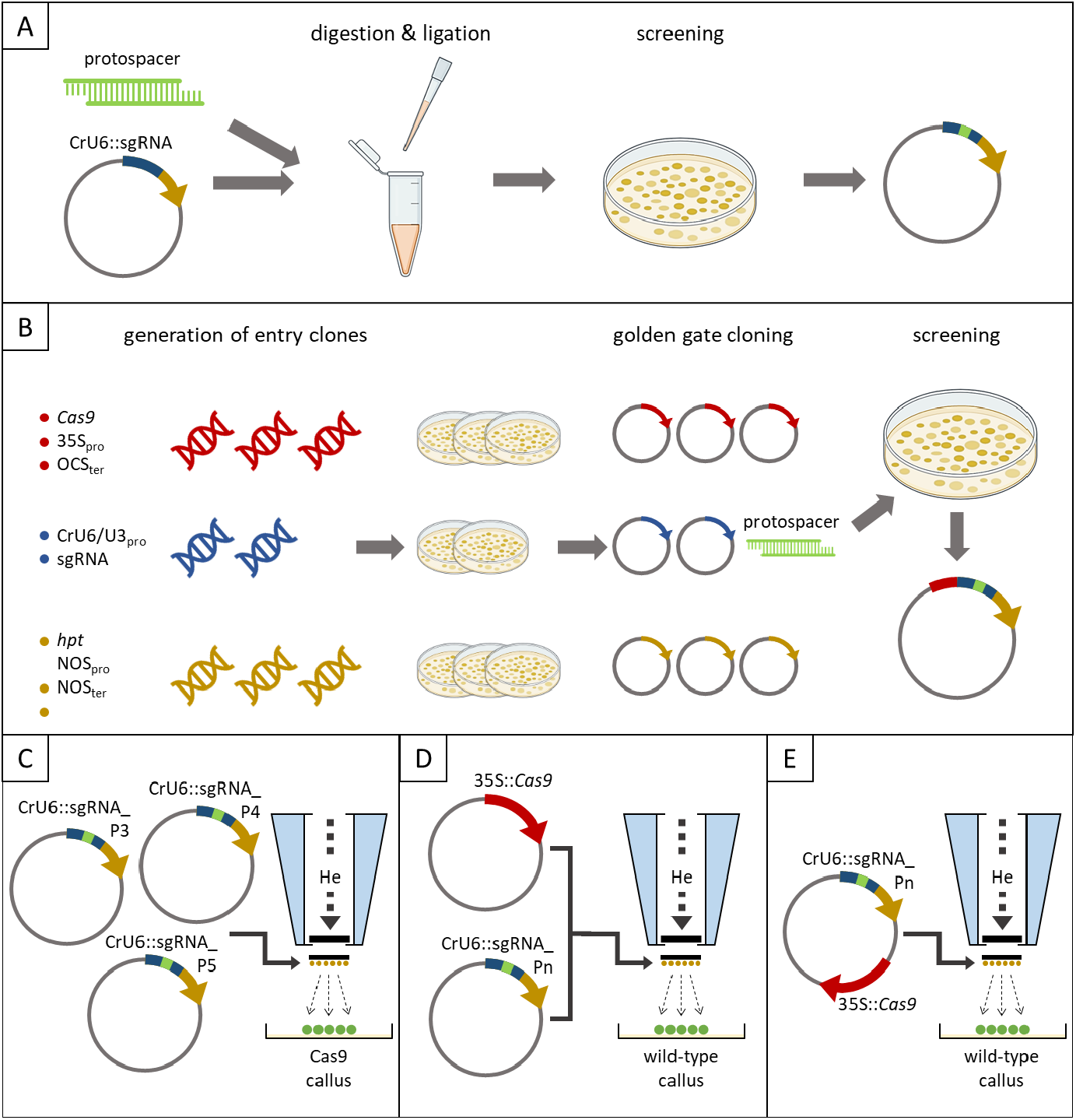
Schematic flow for generation of CRISPR-Cas9 constructs and execution of transformation using biolistic. (A) Simple one tube integration of a protospacer into pre-build sgRNA construct with transcription driven by endogenous CrU6 promoter in contrast to (B) the extensive workflow of generating a construct containing Cas9 driven by a 35S promoter, sgRNA driven by CrU6 promoter and hpt for selection of plants on hygromycin via golden gate cloning. The transformation of Ceratopteris can be done with (C) a mix of sgRNA constructs with Cas9 expressing callus as target, (D) co-transformation of a Cas9 construct with sgRNA constructs using wild-type as target or (E) a construct containing both Cas9 and a sgRNA also using wild-type as target.

Shearing of the transgene can cause a multitude of unwanted side effects like false positives or changes in the phenotype by knockout of endogenous genes. To decrease the effect of shearing, simplify the procedure and highly decrease the labour we created a novel CRISPR-Cas9 system for *Ceratopteris richardii* that is based on the transformation of *Cas9*-expressing mutants with a gene encoding sgRNA. It needs only a construct containing the *Ceratopteris* specific CrU6 promoter, an sgRNA gene with a protospacer, and selection marker reducing the size of the transgenic cassette to ∼2.2 kb. Our system is working in a single tube containing the protospacer dsDNA with matching overhangs and the sgRNA construct (Fig. 2A). The plasmid is first linearized by digesting it with *Bbs*I and then a ligation mix is directly added to the digestion to integrate the protospacer and recyclize the plasmid. We have performed this method with six different protospacers and tested five *E. coli* colonies per protospacer with PCR and digestion with *Bbs*I resulting in a rate 95 % (n = 30) positive integration.

For transformation with a gene gun it is recommended to coat the gold particles with at least 5 μg plasmid DNA. Since the size of the plasmid directly determines the number of molecules per mass means using a smaller plasmid allows for more molecule to bind per gold particle and therefore increasing the efficiency of the transformation. When transforming wild-type as the target for CRISPR-Cas9 the options are to co-transform Cas9 and sgRNA as a binary system (Fig. 2D) or using a single plasmid (Fig. 2E). In a binary system the Cas9-plasmid is much larger than the sgRNA plasmid which means it has to be used in a lower concentration therefore reducing the transformation efficiency. A single plasmid containing all elements in very large in size, allowing only a small number of molecules to bind to the gold particles and therefore reducing the transformation efficiency. Using *Cas9*-expressing callus as target for transformation with sgRNA not only allows for a greater number of molecules to bind to the gold particles and therefore increases the efficiency but also allows a flexibility of easily mixing various protospacers (Fig. 2C), although decreasing their respective concentration.

To generate the *C. richardii* mutant we have built a construct for transformation containing *Cas9* under a CaMV35S promoter and selection of plants on hygromycin B (Fig. 3A). The selection process and regeneration of sporophytes took six weeks. After that the regenerated sporophytes were transferred to soil. We isolated genomic DNA from eleven plants T_0_ plants and performed PCR on full length of *Cas9* that showed 73 % of the plants were positive. We then isolated total RNA and synthesized cDNA from the eight positive plants that showed a transcription of *Cas9* in five plants. One of those five plants was mutant 6 named 35S∷*Cas9*-6 (Fig. 3B) that was also the first to develop spores which is why we decided to continue with this. Five plants of the next generation T_1_ of 35S∷*Cas9*-6 were all tested positive in a PCR on gDNA while only two plants also showed transcription of *Cas9*. One of those two plants was the mutant 4 named 35S∷*Cas9*-6-4 (Fig. 3C). All PCR products were confirmed by Sanger sequencing (Macrogen Europe, Netherlands).

**Figure 3:**
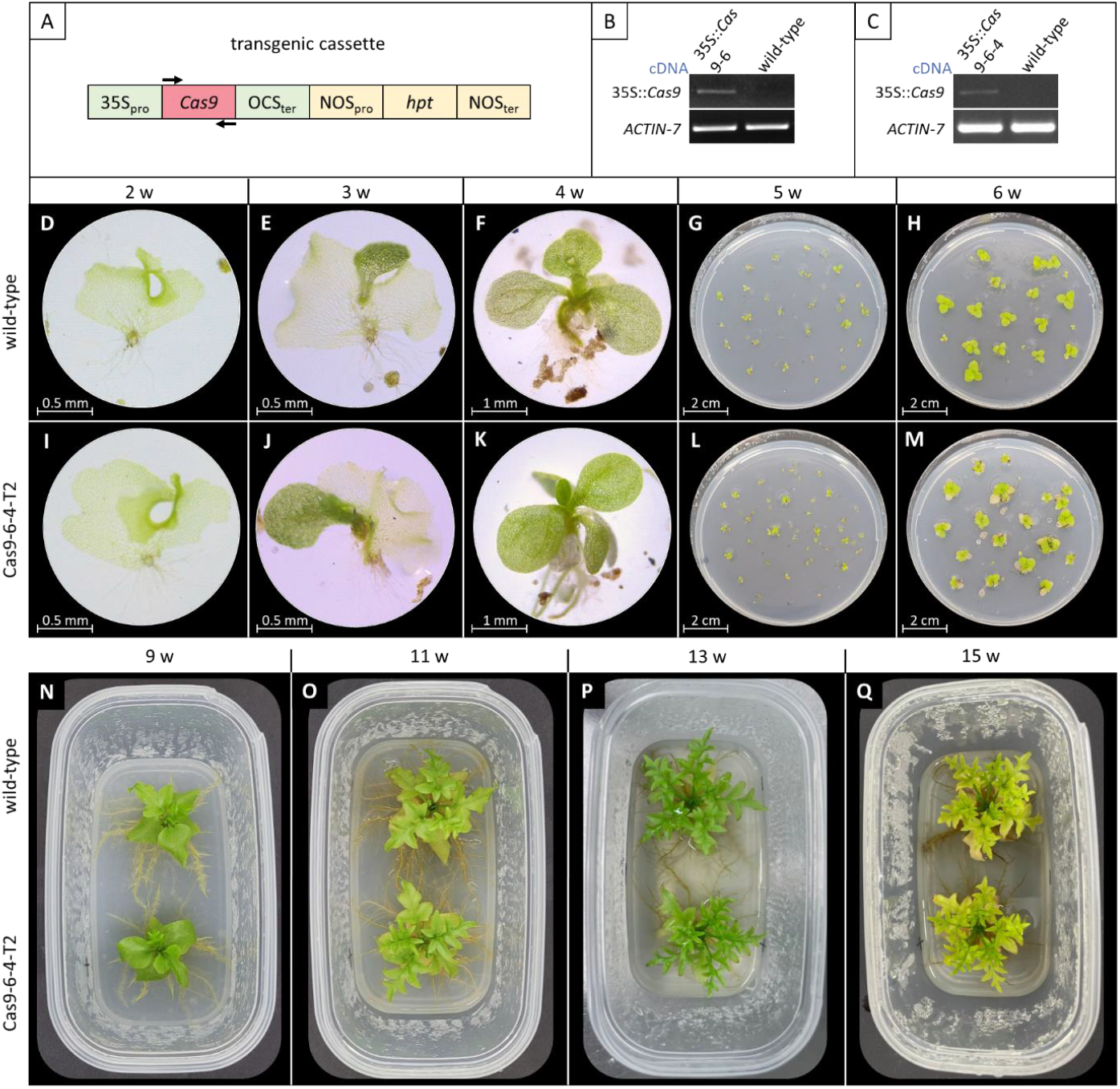
Direct comparison of the phenotype of the Cas9 expressing Ceratopteris richardii mutant line 6-4 with the wild-type covering the entire life-cycle. (A) Cas9 is driven under a CaMV35 promoter and the plants are selected on hygromycin B. (B) cDNA synthesis on total RNA of the Cas9 mutant 6 is positive on the transcription of Cas9. (C) Transcription was confirmed in the T1 generation of mutant 6 (D-H) shows the wild-type and (I-M) the Cas9 mutant development from a hermaphrodite gametophyte to early sporophyte in weekly steps starting with 14 d. (N-Q) shows the wild-type (upper plant) and the Cas9 mutant (lower plant) development from juvenile vegetative to mature reproductive sporophytes.

An important aspect of the *Cas9* expressing mutant line was that it does not produce a mutant phenotype by itself. We used spores harvested from 35S∷*Cas9*-6-4 to demonstrate that the phenotype of the *Cas9* mutant (Fig. 3I-U) is indistinguishable from the wild-type (Fig. 3D-Q) in both gametophyte and sporophyte by comparing the major stages of the life-cycle. The spores of both genotypes were harvested at the same time, dried for two months. Every step was performed simultaneously and under the same conditions for both 35S::*Cas9*-6-4 and wild-type. Pictures of the first four weeks (Fig. 3D-F & I-K) were taken under the binocular Leica ez4w (Leica, Germany). The direct comparison during the first six weeks has shown no noticeable difference in the phenotype despite having the mutant grown on hygromycin selection media. Sterile plastic containers with one nine weeks old sporophyte of 35S::*Cas9*-6-4 and wild-type were prepared to allow for larger growth while remaining under the same conditions. The plants were watered every week with sterile liquid C-fern media diluted 1:1 with sterile ddH_2_O. No hygromycin was added to the media. Pictures were taken up to 15 weeks (Fig 3N-Q). Size of the plants and fronds, colour, shape and count of the fronds as well as root development show no phenotypic differences between the transgenic 35S::*Cas9* and wild-type plants.

## Discussion

Ferns represent an important group of land plants. In contrast to bryophyte model systems such as *Physcomitrium patens* (moss) and *Marchantia polymorpha* (liverwort), and model angiosperms such as *A. thaliana*, rice and many others, little is known about the function of genes in ferns. Rare exceptions are represented by the study of a few mutants obtained e.g. by chemical mutagenesis and gene knock-down harnessing RNAi, the investigation of expression patterns, and the analysis of similarity and homology to genes from other groups of land plants (Eberle et al., 1995; Münster et al., 1997, 2002; Hasebe et al., 1998; Bank, 1994, 1999; Plackett et al., 2018).

The analysis of null mutants can be considered the ,gold standard’ way for defining the function of a gene, because it reveals what happens with an organism if a gene is not functional anymore, indicating what the gene is required for. Exceptions are cases of genetic redundancy, that may require double or even multiple mutants, and cases of lethality, in which conditional mutants might be required to obtain informative clues about gene function.

Functional knock-outs of a protein-encoding gene (null mutants) are relatively easily obtained by generating inserions or deletions (indels) in the CDS of the gene. Even in case of short sequence changes, gene function will be very likely abolished, if the indel is not to close to the 3’ end of the CDS, and if indel length (in nt) is not a number that can be divided by 3 (the length of a codon). In these cases truncated proteins are very likely produced that are not functional due to a translational reading frame shift, or even the mRNA is degraded by NMD.

The CRISPR-Cas9 system provides a relatively easy way to generate short indels in potentially any CDS (Ran et al., 2013; Doudna and Charpentier, 2014). Indeed, the versatility of the sgRNA-mediated CRISPR-Cas9 system for the generation of targeted mutations in plants has already been demonstrated for diverse flowering plant species, including eudicots such as *A. thaliana* and tobacco (*Nicotiana benthamiana*), and monocots such as rice (*Oryza sativa*) and sorghum (*Sorghum bicolor*) (e.g., Jiang et al., 2013). For a single gene the applicability of CRISPR-Cas9 in *C. richardii* has recently been documented (Xiang and Li, 2024). CRISPR-Cas9 is hence an obvious choice for establishing an efficient reverse genetics and genome editing system in *C. richardii*.

Here we report the generation and identification, respectively, of Cas9 expressing plants and CrU6 promoters from *C. richardii*. Using Cas9-expressinfg callus from the transgenic plants for microparticle bombartment introducing DNA-constructs that drive sgRNA under control of a CrU6 promoter willy simplify genome editing in *C. richardii* by CRISPR-Cas9. For example, it will facilitate the introduction of several different sgRNA-encoding constructs to knockout different target genes in parallel. Our system is currently employed to knockout the MIKC^C-^ type MADS-box gene *CRM3* (Münster et al., 1997; Schulz and Theißen, 2025), but might be well-suited in many other projects as well, not necessarily limited to knock-out studies.

## Acknowledgements

Many thanks to Andrew Placket for his valuable help with the establishment of the *Ceratopteris* transformation system. Many thanks also to Annette Becker for numerous valuable discussions on *Ceratopteris* research. In addition we thank also all other members of the ICIPS research unit for fruitful discussions.

